# Development and Testing of New Genetic Markers for the Detection of Invasive Bighead and Silver Carp (*Hypophthalmichthys nobilis* and *H. molitrix*) DNA in environmental water samples from North America

**DOI:** 10.1101/003624

**Authors:** Heather L. Farrington, Christine E. Edwards, Xin Guan, Matthew R. Carr, Kelly Baerwaldt, Richard F. Lance

## Abstract

Invasive Asian bighead and silver carp (*Hypophthalmichthys nobilis* and *H. molitrix*) pose a substantial threat to North American waterways. Recently, environmental DNA (eDNA), the use of species-specific genetic assays to detect the DNA of a particular species in a water sample, has gained recognition as a tool for tracking the invasion front of these species toward the Great Lakes. The goal of this study was to develop new species-specific conventional PCR (cPCR) and quantitative (qPCR) markers for detection of these species in North American waterways. We first generated complete mitochondrial genome sequences from 33 bighead and 29 silver carp individuals collected throughout their introduced range. These sequences were aligned with other common and closely related species to identify potential eDNA markers. We then field-tested these genetic markers for species-specificity and sensitivity in environmental samples. Newly developed markers performed well in field trials, had low false positive rates and had comparable sensitivity compared to current markers. The new markers developed in this study greatly expand the number of species-specific genetic markers available to track the invasion front of bighead and silver carp, and can be used to improve the resolution of these assays. Additionally, the use of the qPCR markers developed in this study may reduce sample processing time and cost of eDNA monitoring for these species.

## Introduction

Invasive aquatic nuisance species pose a major threat to aquatic ecosystems worldwide. In North America, invasive Asian carps, particularly bighead carp (BHC; *Hypophthalmichthys nobilis*) and silver carp (SC; *H. molitrix*), have been very problematic in freshwater ecosystems. Asian carps were imported into the U.S. in the 1970s to control algae in Arkansas fish farms (Freeze and Henderson 1982). Flooding allowed them to escape and establish reproducing populations in the wild by the early 1980s. They have since been steadily dispersing upstream throughout the Mississippi River watershed (Freeze and Hendersen 1982; Tucker et al. 1996). At present, BHC and SC have been found in 23 states, and they have rapidly expanded their population sizes, with BHC and SC representing over 60% of the biomass in some portions of their North American Range (Garvey et al 2012). These filter-feeders cause significant ecological impacts by altering plankton communities at the base of the food chain and outcompeting native species for resources. There is considerable concern that these species will enter the Great Lakes through man-made shipping, sanitation and flood control canals, such as those of the Chicago Area Waterways System (CAWS). Should self-sustaining BHC or SC populations become established in the Great Lakes, these species could potentially cause dramatic ecosystem alterations, leading to negative effects on populations of native fishes and many threatened or endangered plant/animal species (Asian Carp Regional Coordinating Committee 2013). The impact of this invasion on Great Lakes fisheries is of particular concern.

Aquatic organisms shed biological materials (*e.g.*, scales, epithelial cells, slime coats, waste) containing DNA into their environments. This environmental DNA (eDNA) can persist in aquatic environments for extended periods (Dejean et al. 2011, Thomsen et al. 2012), and the eDNA in water samples can be assayed using species-specific genetic markers to determine whether a species of interest may be present. Because eDNA can be detected in water when target species’ populations are at low abundances, eDNA techniques may be particularly helpful in tracking changes in the distributions of aquatic invasive species (Ficetola et al. 2008; Dejean et al. 2012; Jerde et al 2011, Goldberg et al. 2013, Lance and Carr 2012) or identifying locations where threatened or endangered species may occur (Goldberg et al. 2011; Olson et al. 2012; Farrington and Lance, in prep).

Since 2009, eDNA monitoring has been used to track the invasion front of BHC and SC throughout the CAWS, Des Plaines River, and near-shore waters of Lake Michigan. The current eDNA monitoring program employs a single, species-specific genetic marker to detect each species (Jerde *et al*. 2011). The program utilizes conventional polymerase chain reaction (cPCR) analysis, whereby the presence or absence of eDNA is determined by PCR amplification of a target DNA fragment. The PCR-amplified product is then isolated by gel electrophoresis and the DNA is sequenced to confirm the species of origin. The Quality Assurance Project Plan (QAPP) for the Environmental DNA (eDNA) Monitoring of Invasive Asian Carp in the CAWS outlines the detailed procedures for the current planning, collection, filtering and processing of eDNA samples (USACE 2012).

The development of additional BHC and SC eDNA markers could provide a suite of assays to provide multiple lines of evidence or secondary verification for eDNA detections. In addition to cPCR markers, quantitative PCR (qPCR) may be used as an eDNA monitoring tool. The use of qPCR has several potential advantages relative to cPCR, including, typically, more rapid PCR thermal-cycling programs, which can be important for large-scale sampling efforts, a reduced sensitivity in some cases to PCR inhibitors (personal observation; Barnes et al. 2014), and the ability to quantify, to some degree, the amount of DNA in a sample (taking into account inherent variations in DNA extraction recoveries and qPCR-based copy number estimates). Also, while conventional PCR requires specific oligonucleotide binding at *two* locations (the forward and reverse primers) in order to produce a PCR product, hydrolysis probe-based qPCR, which is one of two common qPCR methodologies, may often be a more stringent assay because it requires specific oligonucleotide binding at *three* locations (forward and reverse primers, as well as the internal hydrolysis probe) in order for the reaction to produce a product that emits a fluorescent signal.

Our objectives in this study were to: 1) sequence full mitochondrial (mtDNA) genomes from multiple BHC and SC throughout their North American range to represent the intraspecific genetic variation of each species, 2) use multiple sequence alignments of BHC, SC and other closely related species that may be present in aquatic ecosystems in the Midwestern U.S.A. to design species-specific cPCR and qPCR markers for the detection of BHC and SC in eDNA monitoring programs, and 3) test the specificity and sensitivity of these new markers in detecting BHC and SC in laboratory and eDNA field trials.

## Methods

### Sample Collection, DNA sequencing, and Alignment

Tissue samples (fin clips or livers) were collected from silver and bighead carp populations throughout their introduced range within the Mississippi River watershed (Table 1; Fig 1). Total genomic DNA was extracted using DNeasy Blood and Tissue Kits (QIAGEN Inc.) according to the manufacturer’s instructions. DNA extractions were enriched for mitochondrial DNA using long PCR to amplify the mitochondrial genome as a single 16.6 kb fragment. Primer sequences were S-LA-16S-L 5′-CGATTAAAGTCCTACGTGATCTGAGTTCAG-3′ and S-LA-16S-H 5′-TGCACCATTAGGATGTCCTGATCCAACATC-3′ (Miya and Nishida 2000). QIAGEN LongRange PCR Kit reagents were used to formulate a 25 µL PCR reaction mixture containing 1× LongRange PCR buffer, 500 µM dNTPs, 1.25 U LongRange PCR Enzyme mix, 0.4 µM of each primer, and 1 µL of DNA template. Temperature cycling conditions began with an initial denaturation step of 93°C for 3 min, followed by 10 cycles of 93°C for 15 sec, 62°C for 30 sec and 68°C for 18 min. An additional 29 cycles were then run adding 20 sec to the extension step for each cycle. Because amplification of a single fragment was not successful for all samples (likely due to degraded template DNA), we also attempted to amplify the mitochondrial genome in three shorter, overlapping fragments, using the same PCR chemistry and cycling conditions described above. Primer sequences were designed using Primer3 software (Rozen and Skaletsky 2000) based on BHC and SC complete mitochondrial genome sequences available on GenBank (accession numbers NC_010194, EU343733, JQ231114, HM162839, EU315941, NC_010156). The following primers were designed to amplify fragments of approximately 7.4, 7.0 and 3.0 kb, respectively: LC1-F and R (5′-GAATGGGCTAAACCCCTAAA -3′/ 5′-TCGTAGTGAAAAGGGCAGTC -3′); LC2-F and R (5′-CAGGATTCCACGGACTACAC -3′ / 5′-TTGGGGTTTGACAAGGATAA -3′; LC3-F and R (5′-CATGCCGAGCATTCTTTTAT -3′ / 5′-CAACATCGAGGTCGTAAACC -3′). When agarose gel electrophoresis revealed that all three of the shorter PCR reactions produced bands of the expected sizes, the reaction products were pooled for sequencing.

**Table 1:** Origins of silver and bighead carp samples included in mtDNA genome sequencing and alignment

| **Location** | **Silver Carp (n)** | **Bighead Carp (n)** |
| --- | --- | --- |
| East Lower Mississippi (Yazoo River, Steele Bayou, Big Sunflower River) | 3 | 5 |
| West Lower Mississippi (Red River, Atchafalaya River) | 3 | 6 |
| Arkansas River | 3 | 3 |
| Ohio River (at junction to Mississippi River) | 3 | 3 |
| Mississippi River (Knowlton Lake) | 2 | - |
| Illinois River (LaGrange Reach) | - | 3 |
| Illinois River (Marseilles Reach) | 3 | 3 |
| Illinois River (Starved Rock) | 2 | 3 |
| Mississippi River (Laketon, KY) | 3 | 1 |
| Upper Mississippi River (Pool 20) | 1 | 3 |
| Upper Mississippi River (Pool 26) | 3 | 3 |
| Missouri River (north of Omaha) | 3 | - |

**Figure 1:**
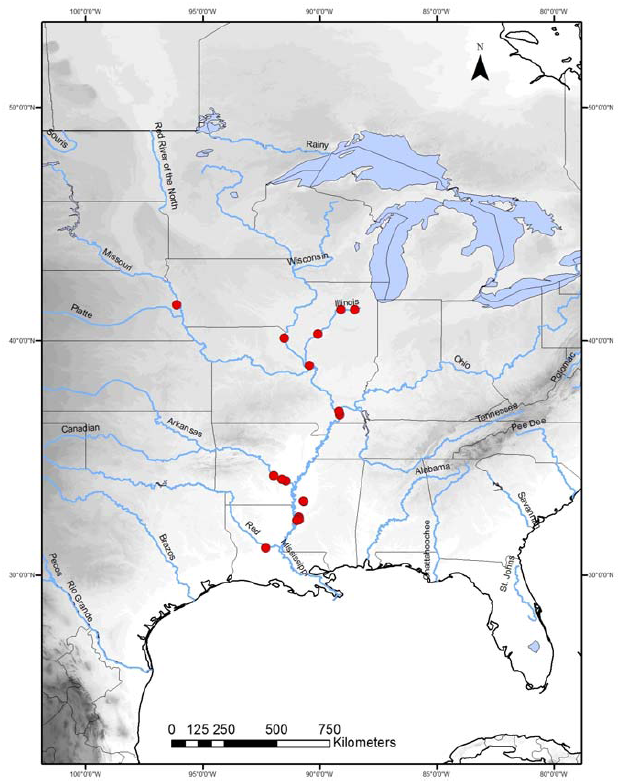
Geographic distribution of sample collection for mitochondrial DNA sequencing

PCR products were purified using ZR-96 DNA Clean and Concentrator-5 kits (Zymo Research) and prepared for next-generation sequencing using Nextera DNA Sample Preparation Kits (Illumina, Inc.); Nextera Index Kits were used to pool up to 96 libraries into a run for sequencing. Sequencing was performed on the Illumina MiSeq system, using 150 bp paired-end reads. MiSeq Reporter Software was used to sort the resulting pool of sequences by the indices to identify the sequences arising from each sample. Mitochondrial genomes were assembled by aligning the reads of each individual to a reference sequence of the appropriate species from GenBank (see above for accession numbers) using Geneious software v.6 (Biomatters Ltd., Auckland, New Zealand). Consensus sequences generated for each individual were exported and aligned, along with sequences of some related cyprinid fish species that may be present in the same North American regions as BHC and SC (common carp, grass carp and black carp; GenBank accession numbers NC_010288.1, NC_018035.1, NC_018039.1, NC_018036.1, NC_011141.1). Alignments were carried out using the default settings in MUSCLE (Edgar 2004) as implemented in Geneious V6.

### Marker Design

Marker loci were designed using the multiple sequence alignment of complete mitochondrial genomes of bighead carp, silver carp, and several related species (listed above). Potential PCR primer sites were chosen by identifying sequence regions that demonstrated no mismatches within the target taxa and that maximized differences between target and non-target taxa. Because eDNA may experience rapid degradation by environmental conditions, marker loci were designed to be short (<400 bp) to increase amplification probability. Primer3 (Rozen and Skaletsky 2000) was used to design cPCR primers and qPCR primer/probe sets with preference for primers that contained 3'-end mismatches to homologous DNA in non-target species. All qPCR probes were labeled with 6FAM as the 5' fluorescent tag, and TAMRA as the 3' quencher. Due to the limited genetic divergence between bighead and silver carp, we also developed a series of general BHC/SC markers that may detect both species.

### Marker testing for specificity and efficacy in eDNA field trials

Unless otherwise noted, newly designed cPCR markers were tested using 25 µL reactions containing 1× Platinum^®^ *Taq* PCR buffer (Invitrogen), 200 µM dNTPs, 1.5 mM MgCl_2_, 0.2 µM of each primer, 1.25 U Platinum^®^ *Taq* polymerase (Invitrogen), and 1 µL DNA template. Temperature cycling conditions began with an initial denaturation step of 94°C for 10 min, followed by 45 cycles of 94°C for 1 min, 52°C for 1 min and 72°C for 1 min 30 sec, with a final elongation step at 72°C for 7 min. Amplification products of cPCR assays were purified using E-Gel SizeSelect Gels (Life Technologies) and sequenced using an ABI 3500XL Genetic Analyzer with BigDye chemistry and standard sequencing protocols. Resulting sequences were compared against BHC and SC reference DNA sequences and subjected to GenBank BLAST searches to identify the source species of the amplification product.

All qPCR reactions were run in 20µl volumes containing 1X TaqMan^®^ Environmental Master Mix, 0.54µM of each primer, 0.125 µM of the probe, and 1 µL of DNA template. Temperature cycling began with an initial denaturation step at 95°C for 10min, followed by 40 cycles of 95°C for 15 sec and 60°C for 1 min. qPCR reactions were run on a ViiA™ 7 Real-Time PCR System (Applied Biosystems). qPCR reactions were considered positive if the amplification curve crossed the fluorescence detection threshold by the end of the 40 cycle qPCR run.

Both cPCR and qPCR markers were tested for: 1) species-specificity, 2) ability to amplify target species DNA from eDNA samples collected in areas of known BHC and SC presence, 3) false-positive amplification from eDNA samples that likely do not contain Asian carp DNA, and 4) limits of detection, or sensitivity, in targets species (*i.e.*, the minimum amount of starting DNA that can result in a detectable cPCR or qPCR product).

Species-specificity of both cPCR and qPCR assays was tested using a panel of genomic DNA (1 ng/µL) from individuals of the target species and 29 additional species likely to be present in the CAWS. This panel included closely related, non-target species such as shiners, common carp, goldfish, and grass carp (Table 2). If cPCR markers amplified non-target species, annealing temperatures were adjusted in an attempt to eliminate non-target amplification. The cPCR and qPCR markers that amplified the target species and showed little cross-amplification in non-target species were further tested using field-collected water samples from Steele Bayou, a backwater flood control area near the Yazoo River’s confluence with the Mississippi River near Vicksburg, MS. U.S.A. Steele Bayou is locally known to have well-established BHC and SC populations with high densities (pers. commun., A. Katzenmeyer). We also tested for amplification of BHC and SC in water samples collected from a small tributary of Fishing Creek (Clinton County, PA, U.S.A), an area outside the introduced range of BHC and SC. These samples have all the typical components of environmental water samples, but were free of target DNA, and thus provided test cases to detect potential non-target amplification within naturally occurring DNA pools. In all cases, surface water samples were collected in 50 mL conical tubes. In the laboratory, tubes were centrifuged at maximum speed (4000 g) for 30 min at 4° C. The supernatant was poured off and DNA was extracted from the remaining pellet of material using a modified cetyltrimethyl ammonium bromide (CTAB)/chloroform protocol (Doyle and Doyle 1987). Each cPCR and qPCR marker was tested on a panel of 44 Steele Bayou and 44 Fishing Creek samples, with 4x replication of PCR reactions to evaluate the detection rate of these species from areas of known presence and the potential false positive rate from waters where they are absent. The performance of all new markers, as measured by rate of detection, was compared to the cPCR markers for BHC and SC from Jerde et al. 2012 (primers HN203-F & HN498-R and HMF-2 & HMR-2, respectively), which are currently used in the Asian carp monitoring QAPP (USACE 2012). We refer to these markers here as QAPP-SC and QAPP-BHC.

**Table 2:**
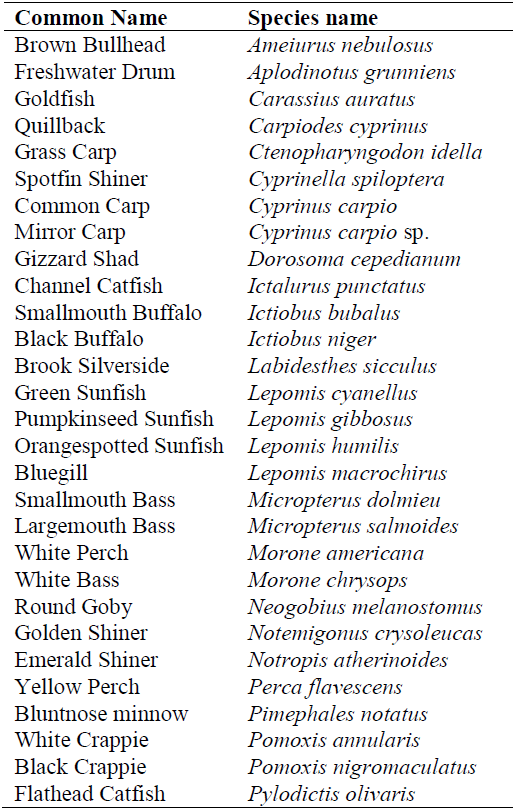
Panel of 29 non-target fish species collected from the CAWS used for testing of primers for cross-species amplification.

Markers with high detection rates and low false-positive rates in environmental samples were subjected to sensitivity testing. Genomic DNA of SC and BHC was extracted and the concentration of each was normalized to 1 ng/µL. A serial 1:10 dilution series was prepared and markers were tested across the concentration range of 0.1 ng/µL (10^-1^) through 10^-7^, with four replicate cPCRs or qPCRs at each concentration. A limitation to the use of genomic DNA in sensitivity testing for cPCR markers is that the number of marker copies present in the normalized DNA extractions, and therefore available for PCR amplification, is unknown. To estimate starting copy number in qPCR reactions, each qPCR marker was cloned into a bacterial plasmid vector using TOPO^®^ Cloning kits (Life Technologies) as per the manufacturer’s instructions. Successfully cloned bacterial colonies were cultured and plasmids extracted using Qiagen Miniprep plasmid extraction kits. The estimated number of plasmids in the resulting elutions was calculated using the combined base pair length of the plasmid and marker insert, a standard DNA base-to-Daltons conversion for double-stranded DNA (650 Daltons/base; Roche Applied Science 2011), a Daltons-to-nanograms conversion, and DNA mass quantification of elutions using a NanoDrop 1000. A dilution series of the plasmid elution was then used to generate a standard curve for estimation of copy number in the qPCR reactions.

To test whether the throughput of eDNA screening methods could be increased by assaying for multiple markers simultaneously within single PCRs, several markers were combined in pairs for multiplex cPCR or qPCR reactions. For cPCR, the QAPP-SC and SC-1 markers were combined. For qPCR, three primer sets were tested: BH-TM1/BH-TM2, SC-TM4/SC-TM5, and AC-TM1/AC-TM3. For qPCR multiplexing, the two markers utilized probes with different fluorescent labels (FAM or VIC). A genomic DNA dilution series and plasmid standards were again used for testing, with markers and standards run both individually and in combination in order to directly compare sensitivity in single versus multiplex reactions. qPCR reactions were prepared as described above, with both sets of primers and probes added to the reaction, and the same temperature cycling conditions.

## Results

We generated complete mtDNA sequences for 33 BHC and 29 SC individuals (Table 1); all DNA sequences were submitted to GenBank (Accession numbers XXXX to XXXX). Average whole genome sequence coverage for the 62 haplotypes sequenced was 1595X (range 3-11971X coverage). Total length of the aligned BHC and SC genomes was 16620 bp. There was very little sequence variation within species, with only 40 (0.24%) and 34 (0.20%) variable sites for BHC and SC, respectively. When species alignments were combined, there were a total of 823 (4.95%) variable sites across the mitochondrial genome. These genomes were aligned with mtDNA genomes from closely related species obtained from GenBank, including common, grass and black carp for identification of potential species-specific eDNA markers.

Based on the alignment of mitochondrial genomes, we initially designed 12 SC, 11 BHC and 16 general BHC/SC cPCR markers. For TaqMan^®^ qPCR, we initially designed five markers for BH, six for SC, and three general BHC/SC markers. Based on results from the initial cross-species screening, several markers amplified non-target species and were not further investigated, reducing the number of potential markers for testing to six for cPCR and eight for qPCR (Table 3). We focused all subsequent field and sensitivity testing on the markers with high affinity for the target species and little or no amplification of other species.

**Table 3.**
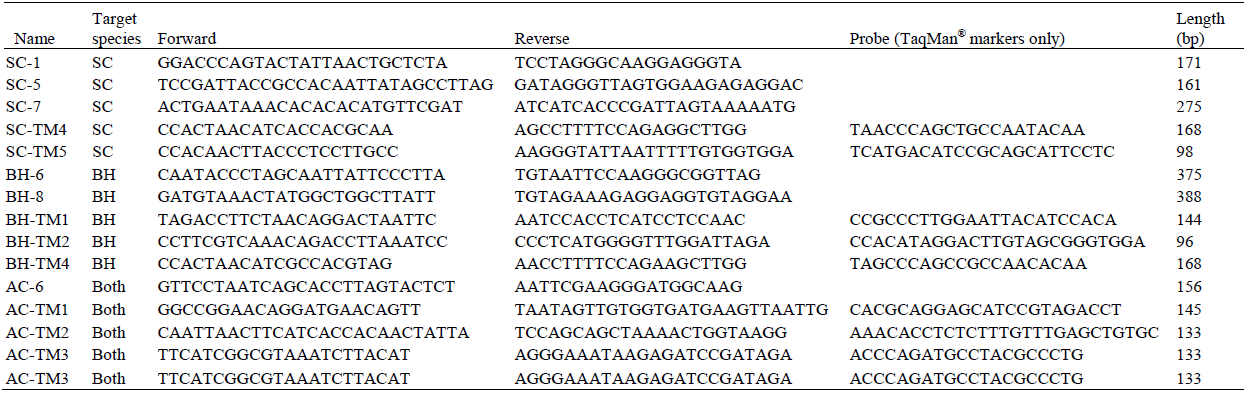
Primer sequences of markers used for field and sensitivity testing. For targeted species, BH=bighead carp, SC=silver carp, Both=primers that could potentially be used for non-specific detection of both BHC and SC. Names containing TM indicates TaqMan^®^ qPCR markers.

**Table 4:** Number of positive detections noted for each marker tested using 44 eDNA field samples. Steele Bayou samples were collected from an area of high concentrations of both BHC and SC, whereas Fishing Creek samples were collected from an area where carp are absent. QAPP-SC and QAPP-BH are the markers currently used for eDNA testing. Names containing TM indicate TaqMan^®^ qPCR markers.

| Marker | Steele Bayou | Fishing Creek |
| --- | --- | --- |
| **Silver Carp:** |  |  |
| QAPP-SC | 28 | 0 |
| SC-1 | 32 | 0 |
| SC-5 | 25 | 0 |
| SC-7 | 23 | 0 |
| SC-TM4 | 26 | 0 |
| SC-TM5 | 32 | 0 |
| **Bighead Carp:** |  |  |
| QAPP-BH | 0 | Not tested |
| BH-6 | 6 | 0 |
| BH-8 | 6 | 0 |
| BH-TM1 | 9 | 0 |
| BH-TM2 | 7 | 0 |
| BH-TM4 | 6 | 0 |
| **Bighead and Silver Carp:** |  |  |
| AC-6 | 30 | 0 |
| AC-TM1 | 25 | 0 |
| AC-TM2 | 30 | 0 |
| AC-TM3 | 28* | 0 |

\* - Tested with only 42 samples

Assays of the 44 Steele Bayou samples with the established markers QAPP-SC and QAPP-BHC resulted in 28 (64%) positive SC detections and 0 (0%) positive BHC detections. All of the newly designed BHC markers performed better than the QAPP-BH marker, with the highest detection rate from the qPCR marker BH-TM2 (9 of 44, 20%). In comparison to the QAPP-SC marker, all newly-designed SC markers had similar or higher numbers of detections, with the highest detection rates from cPCR marker SC-1 and qPCR maker SC-TM5, both with 32 positive samples (73%). Positive detection rates were 57-68% for the general (BHC/SC) markers. For the Fishing Creek samples, none of the new cPCR markers produced bands in the same size range as target species and none of the qPCR markers produced quantifiable fluorescence.

All the tested markers consistently yielded positive results from genomic DNA down to at least the 10^-3^ dilution (0.001 ng/µL). Three SC (SC-5, SC-7 and SC-TM4) and one BHC/SC marker (AC-TM2) had consistent detections at 10^-4^, and nearly all markers had >50% detection rates among the four replicates at the 10^-4^ dilution level, which is estimated to have copy numbers in the single digits (Table 5). Detections became more stochastic at concentrations below 10^-4^, as expected for samples with extremely low copy number (average of ≤1 marker copy per reaction). Sensitivity of the new BHC and SC markers was comparable to the QAPP markers.

**Table 5:** Sensitivity testing. Estimated marker copy numbers per dilution are based on averages calculated across all replicates of qPCR sensitivity trials using a plasmid DNA standard. AC markers were tested using SC dilutions. The number of positive detections out of four replicates is noted for each marker and dilution level. U=Undetermined copy number. Amplifications at these levels are likely due to stochasticity of PCR at such low DNA concentrations.

| Dilution: | $10^{-1}$ | $10^{-2}$ | $10^{-3}$ | $10^{-4}$ | $10^{-5}$ | $10^{-6}$ | $10^{-7}$ |
| --- | --- | --- | --- | --- | --- | --- | --- |
| **Silver Carp:** |  |  |  |  |  |  |  |
| Estimated Copy Number: | 6051 | 508 | 46 | 3.4 | 1.6 | U | U |
| QAPP-SC | 4 | 4 | 4 | 4 | 0 | 0 | 0 |
| SC-1 | 4 | 4 | 4 | 3 | 0 | 0 | 0 |
| SC-5 | 4 | 4 | 4 | 4 | 0 | 2 | 0 |
| SC-7 | 4 | 4 | 4 | 4 | 0 | 0 | 0 |
| SC-TM-4 | 4 | 4 | 4 | 4 | 1 | 1 | 0 |
| SC-TM-5 | 4 | 4 | 4 | 3 | 2 | 1 | 1 |
| **Bighead Carp:** |  |  |  |  |  |  |  |
| Estimated Copy Number: | 2193 | 230 | 16 | 2.2 | 1.1 | U | U |
| QAPP-BH | 4 | 4 | 4 | 4 | 1 | 1 | 0 |
| BH-6 | 4 | 4 | 4 | 1 | 0 | 0 | 0 |
| BH-8 | 4 | 4 | 4 | 3 | 0 | 1 | 0 |
| BH-TM-1 | 4 | 4 | 4 | 3 | 1 | 0 | 0 |
| BH-TM-2 | 4 | 4 | 4 | 3 | 0 | 0 | 0 |
| BH-TM-4 | 4 | 4 | 4 | 3 | 0 | 1 | 0 |
| **Bighead and Silver Carp:** |  |  |  |  |  |  |  |
| Estimated Copy Number: | 6051 | 508 | 46 | 3.4 | 1.6 | U | U |
| AC-6 | 4 | 4 | 4 | 2 | 0 | 1 | 0 |
| AC-TM-1 | 4 | 4 | 4 | 2 | 2 | 0 | 0 |
| AC-TM-2 | 4 | 4 | 4 | 4 | 2 | 0 | 0 |
| AC-TM-3 | 4 | 4 | 4 | 3 | 0 | 0 | 0 |

Multiplexing of cPCR markers was found to be unfeasible for high throughput processing and analysis using standard gel electrophoresis equipment in our lab. The new cPCR markers were all designed to be in the size range of 200-300 base pairs to increase the potential for amplification of degraded eDNA, therefore, amplicon base pair lengths were too similar for clear differentiation of bands on 2% agarose gels; gel-based isolation of fragments for sequencing would also be infeasible with this combination of amplicon lengths and electrophoresis equipment. Longer-running or higher density gels may have allowed more reliable separation of different cPCR marker bands but may somewhat negate the cost and time savings gained from multiplexing. Trials of multiplexed qPCR markers were successful for all combinations of markers tested, with no substantial reduction in marker sensitivity (limits of detection; Table 6).

**Table 6:** Multiplexing of qPCR markers. Average Ct values (copy numbers) across 24 replicates for markers run individually and combined in a multiplex reaction.

| Dilution: | 10<sup>-2</sup> | 10<sup>-3</sup> | 10<sup>-4</sup> | 10<sup>-5</sup> |
| --- | --- | --- | --- | --- |
| **Silver Carp:** |  |  |  |  |
| SC-TM4 | 30.6 (195-424) | 34.1 (10-50) | 37.3 (0-12) | 38.7 (0-2.4) |
| SC-TM5 | 30.4 (872-1498) | 33.7 (60-208) | 37.4 (0-30) | 38.1 (0-11) |
| Combined: |  |  |  |  |
| SC-TM4 | 30.7 (203-422) | 34.2 (11-50) | 37.9 (0-5) | 38.4 (0-3) |
| SC-TM5 | 30.4 (1145-1792) | 33.9 (81-200) | 37.5 (2-23) | 38.4 (0-14) |
| **Bighead Carp:** |  |  |  |  |
| BH-TM1 | 31.2 (107-244) | 34.6 (8-28) | 38.2 (0-4) | 39.0 (0-1) |
| BH-TM2 | 31.2 (120-216) | 34.8 (7-19) | 37.8 (0-4) | 39.3 (0-1) |
| Combined: |  |  |  |  |
| BH-TM1 | 30.6 (118-263) | 34.3 (8-23) | 37.3 (0-5) | 38.3 (0-1) |
| BH-TM2 | 31.1 (137-201) | 34.6 (12-24) | 38.1 (0-7) | 38.7 (0-1) |
| **Bighead and Silver Carp:** |  |  |  |  |
| AC-TM1 | 31.1 (241-358) | 34.5 (16-65) | 38.0 (0-12) | 38.9 (0-3) |
| AC-TM3 | 29.1 (225-398) | 32.3 (16-52) | 36.0 (0-6) | 37.7 (0-1) |
| Combined: |  |  |  |  |
| AC-TM1 | 31.1 (283-536) | 34.5 (22-91) | 37.7 (0-16) | 39.4 (0-1) |
| AC-TM3 | 29.1 (252-501) | 32.4 (19-80) | 35.9 (0-12) | 37.6 (0-2) |

## Discussion

The large number of mtDNA haplotypes generated in this study allowed us to capture inter- and intra-species genetic variation in SC and BHC across their introduced, North American range. This information, along with comparisons to DNA sequences from related species found in the central United States, aided in the design of cPCR and qPCR markers specifically for eDNA testing for SC and BHC in their introduced range. Effective design of PCR-based assays for the differential or discriminatory detection of species requires that sequence differences among taxa be clustered so that multiple differences among taxa are grouped into the length of a PCR primer and two or more of these areas are grouped within a few hundred base pairs. Because SC and BHC are closely related and have very low levels of sequence divergence across their mitochondrial genomes, a very limited number of sites demonstrated a sufficient number of clustered polymorphism to develop effective species-specific markers. Despite careful selection of markers to maximize differences among species, cross-amplification was observed in at least one non-target species for many markers, resulting in the elimination of nearly 70% of the originally designed markers. Despite these difficulties, we were able to design multiple cPCR and qPCR markers that specifically detect SC and BHC in field-collected water samples in North America.

In field trials, the new species-specific markers developed in this study generally had detection rates similar to or higher than the markers currently used to detect the presence of BHC and SC DNA in environmental water samples, with similar levels of sensitivity at low concentrations of target DNA. Further, confounding or efficiency-diminishing factors (e.g., amplicons that result in gel bands of similar size to those obtained for the target species or nontarget fluorescence in qPCR trials) were not observed, indicating that these markers would be suitable as a high-throughput assays to detect the presence of BHC and SC from environmental water samples. Multiplexing of qPCR markers was successful in genomic DNA trials, suggesting that multiplexing may be feasible in eDNA screening, increasing throughput of the assays. However, performance of multiplexing reactions with field eDNA samples remains to be tested, and additional combinations of the various markers could be employed following further testing.

In addition to potential improvements in sensitivity and throughput by the markers developed in this study, the availability of multiple new cPCR markers for eDNA screening of SC and BHC in water samples may help increase the accuracy of eDNA monitoring programs. eDNA samples are largely comprised of randomly fragmented, low-abundance DNA targets. The current program uses a single marker locus to detect the presence of SC or BHC in environmental samples, which may be sensitive to random degradation of the single marker. The use of multiple marker loci would improve overall detection rates and provide stronger evidence for the presence of BHC and SC DNA in the water. Further, the addition of qPCR technology to eDNA screening provides the transition from simple presence/absence data provided by cPCR to the generation of data related to DNA concentration in field samples. This additional information *may* help estimate *relative* abundance or biomass of species of interest in the sampled waterway. qPCR may also reduce sample screening time by eliminating the need for gel electrophoresis and sequence verification. The new qPCR and cPCR markers developed in this study therefore represent a significant expansion of the tools available to detect the invasion of SC and BHC in North America and may improve the accuracy, resolution, and throughput of eDNA monitoring programs for these species in the future.

## Acknowledgements

We thank James Lamer, Meredith Bartron, Jack Kilgore, Steven George, and Alan Katzmeyer for providing tissue or eDNA samples; Kelly Baerwaldt, Denise Lindsay, and Marianne Hynum for logistical support; Michael Jung and Karen Bascom for assistance with lab work; Emy Monroe for reviewing this manuscript prior to submission; and Burgund Bassüner for assisting with preparation of the map of the collection sites. Funding was provided by the Asian Carp Regional Coordinating Committee.

## Author Contributions

H.L.F., C.E.E., R.F.L. designed study, performed sequence alignments, designed primers fro new markers. H.L.F., X.G., M.R.C. performed fieldwork and laboratory testing. K.B. participated in study design, sample collection, and project management. All authors participated in preparation of manuscript.

## Data Accessibility

DNA Sequences: Genbank accessions XXXX-XXXX

